# Control of Epithelial Tissue Organization by mRNA Localization

**DOI:** 10.1101/2024.12.02.626432

**Authors:** Devon E. Mason, Thomas D. Madsen, Alexander N. Gasparski, Neal Jiwnani, Terry Lechler, Roberto Weigert, Ramiro Iglesias-Bartolome, Stavroula Mili

## Abstract

mRNA localization to specific subcellular regions is common in mammalian cells but poorly understood in terms of its physiological roles^1–6,7^. This study demonstrates the functional importance of *Net1* mRNA, which we find prominently localized at the dermal-epidermal junction (DEJ) in stratified squamous epithelia. *Net1* mRNA accumulates at DEJ protrusion-like structures that interact with the basement membrane and connect to a mechanosensitive network of microfibrils. Disrupting *Net1* mRNA localization in mouse epithelium alters DEJ morphology and keratinocyte-matrix connections, affecting tissue homeostasis. mRNA localization dictates Net1 protein distribution and its function as a RhoA GTPase exchange factor (GEF). Altered RhoA activity is in turn sufficient to alter the ultrastructure of the DEJ. This study provides a high-resolution *in vivo* view of mRNA targeting in a physiological context. It further demonstrates how the subcellular localization of a single mRNA can significantly influence mammalian epithelial tissue organization, thus revealing an unappreciated level of post-transcriptional regulation that controls tissue physiology.

## Main

mRNA localization has emerged as a prevalent level of post-transcriptional regulation not only in model systems but also in higher organisms. Indeed, in diverse mammalian cell types, a large fraction of the transcriptome is targeted to specific subcellular destinations and adopts distinct distribution patterns^1–7^. Nevertheless, whether the localization of individual mRNAs is important for mammalian physiology remains poorly characterized, especially in tissues outside the nervous system^8,9^.

A well-studied localization pathway targets mRNAs to peripheral protrusions of mammalian mesenchymal cells through active kinesin-dependent trafficking on microtubules^7,10–12^. Localization to protrusions of mRNAs, such as *NET1* and *RAB13*, plays an important role in mesenchymal migration in cultured cell systems^13–16^. Mechanistically, mRNA location specifies the site of protein synthesis, which can influence the function of the encoded polypeptide. Specifically, mRNA targeting to particular local micro-environments promotes certain co-translational interactions of the nascent protein thus guiding binding partner selection^13,15,17^. In the case of NET1, localization of *NET1* mRNA at protrusions promotes NET1 protein association with a membrane-bound scaffold^15^. Plasma membrane-associated NET1 acts as a guanine nucleotide exchange factor (GEF) for the small GTPase RhoA, a central regulator of the cytoskeleton^18,19^. In contrast, perinuclear *NET1* mRNA promotes nascent NET1 binding to importins leading to its nuclear sequestration. In this way, the location of the *NET1* mRNA modulates cytoskeletal dynamics and influences complex cellular processes like mesenchymal cell migration *in vitro*^15^. Protrusion-localized mRNAs, including *NET1*, have also been observed at the basal surface of *in vitro* cultured epithelial cells as well as in intestinal enterocytes *in vivo*^1,2^. However, the functional significance of this epithelial basal localization is unknown.

Here we examine the functional role of protrusion mRNA localization in mouse epithelial physiology. We find that, strikingly, protrusion-localized mRNAs are targeted to uncharacterized protrusion-like structures formed by basal keratinocytes in mammalian stratified epithelial tissue. These formations at the dermal-epidermal junction (DEJ) are connected to mechanosensitive microfibrils in the extracellular matrix. Using specific, sequence-blocking anti-sense oligos to prevent *Net1* mRNA localization to the DEJ, we show that *Net1* mRNA location is necessary for maintaining DEJ architecture and epithelial homeostasis through the RhoA pathway. This study connects mRNA localization to unexplored aspects of epithelial physiology and highlights this prevalent level of post-transcriptional regulation as an important contributor to tissue function in higher organisms.

## Results

### Basal accumulation of Net1 mRNA is conserved across epithelial tissues

To probe the function of protrusion-localized mRNAs in epithelia *in vivo*, we first surveyed the localization of several protrusion localized mRNAs^11,20^ (i.e. *Net1, Cyb5r3, Palld, Kif1c, Pkp4*) across various mouse epithelial tissues that are architecturally distinct. These tissues included monolayered tubular epithelial structures (intestine and kidney) as well as stratified squamous epithelia (tongue and skin) (Fig. 1a). To allow spatial quantitative assessments, RNA detection was accompanied by staining with a combination of wheat germ agglutin (WGA) and β-catenin to delineate overall tissue architecture and cell borders (Fig. 1b and Extended Data Fig. 1a,b). To permit comparison across tissues, we focused in each case on the basal layer of cells in contact with the extracellular basement membrane (BM). The BM was demarcated by WGA providing a comparable basal boundary between BM-associated cells in different tissues. Tissues were manually segmented as individual layers (Fig. 1a,b and Extended Data Fig. 1a,b; BM marked by a yellow dashed line) and RNA signal was quantified along the apico-basal axis (Fig. 1c). We define the RNA amount in the 30% basal part of this epithelial layer as the ‘basal mRNA fraction’ (Fig. 1c and Extended Data Fig. 1a). Interestingly, most of the tested mRNAs accumulate in the basal part of the epithelial layer, as opposed to the non-targeted control *Gapdh* mRNA (Fig. 1d and Extended Data Fig. 1c). There was also evident variability in basal mRNA targeting both among tissues and between protrusion-localized mRNAs, suggesting transcript- and tissue-specific differences in the magnitude of mRNA localization. Nevertheless, mRNAs like *Net1* and *Cyb5r3* emerged as consistently basally localized mRNAs in all tissues tested.

**Fig. 1:**
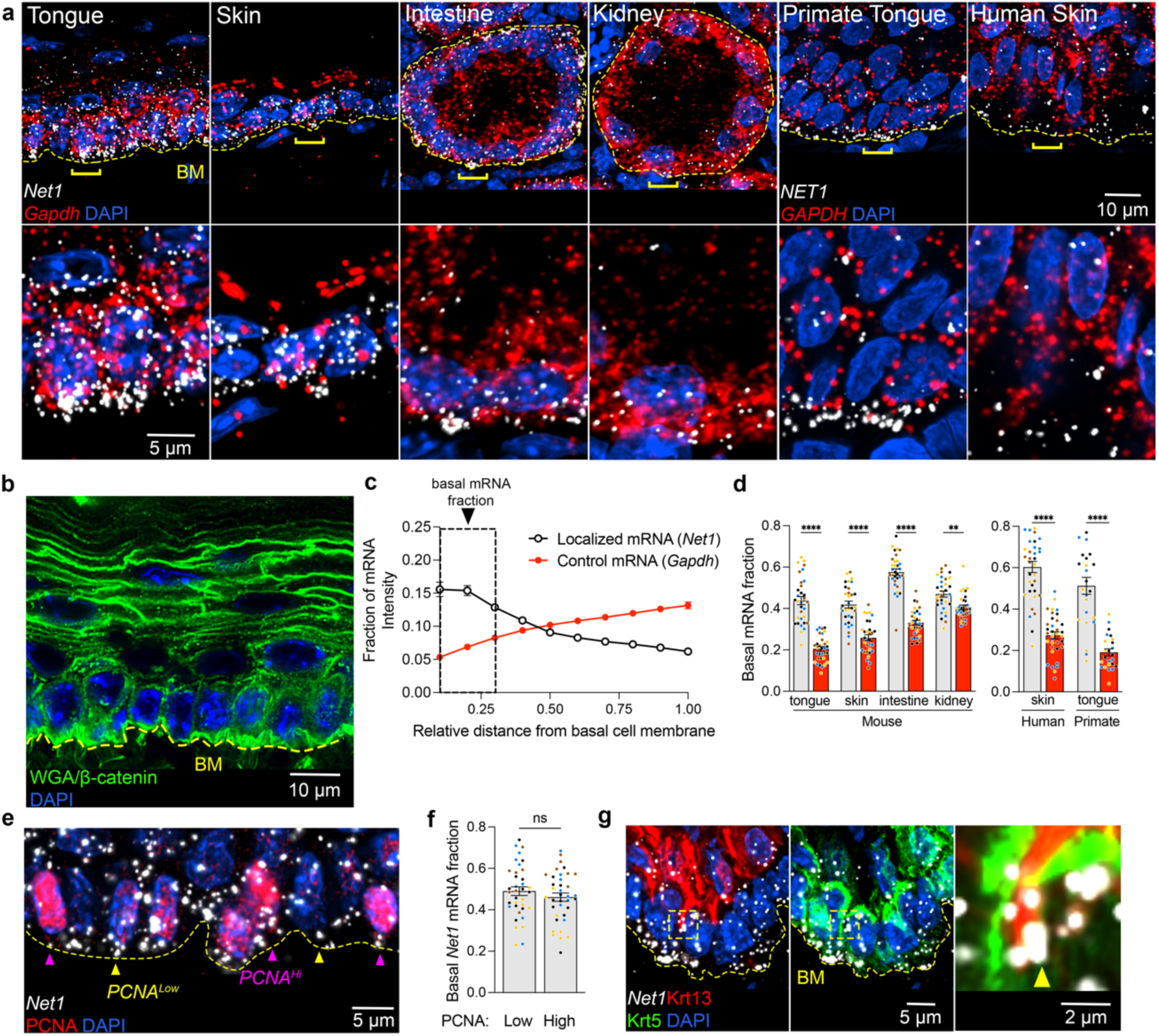
Conserved basal localization of *Net1* mRNA across epithelial architectures and cellular contexts. **a**, Upper panels: Representative images of *Net1* and *Gapdh* mRNAs in mouse tongue, skin, intestine, and kidney as well as images of primate tongue and human skin. Dashed yellow line indicates the basement membrane (BM). Bottom panels: zoomed in regions around areas indicated by brackets. **b**, Representative image of wheat germ agglutin (WGA) and β-catenin, as well as nuclei by DAPI in mouse tongue. Staining was used for segmentation of tissue layer boundaries. **c**, *Net1* and *Gapdh* fluorescent intensity in subsections starting at the BM to the most apical portion of the cell layer. Boxed region indicates RNA amount defined as the basal mRNA fraction. **d**, *Net1 mRNA* (white bars) exhibits basal enrichment relative to *Gapdh* (red bars) across tissues and species. **e**, Visualization of *Net1* localization and PCNA in mouse tongue. Yellow arrowheads indicate PCNA^Low^ and pink arrowheads indicate PCNA^Hi^ cells. **f**, Basal fraction of *Net1* in PCNA^Hi^ vs PCNA^Lo^ cells. **g**, Representative images of basal (Krt5+) and spinous (Krt13+) cells with an inset showing *Net1* mRNA localized to the basal tips of Krt13+ cells (yellow arrowhead). Bar graph data are measurements from individual tissue regions (ROIs) with mean ± SEM; n = 31 (d) or 36 (f) ROIs from N = 3-4 mice. Data from individual mice are represented with different colors. ** P ≤ 0.01, **** P ≤ 0.0001, ns: non-significant by Brown-Forsythe and Welch ANOVA followed by Dunnett’s T3 multiple comparison test (d), or unpaired two-tailed Student’s *t*-test (f).

To look further into these candidates, we surveyed published RNA-seq and tissue RNA imaging datasets. Interestingly, *Net1* is specifically enriched in basal cells of the skin and tongue; *Net1* is also nearly 4-times as abundant as the 5 next most highly expressed Rho-family GEFs (Extended Data Fig. 1d, e)^21–23^. Furthermore, *Net1* expression is 9-times higher in keratinocytes than dermal fibroblasts of the skin pointing to a potential important function in the epidermal compartment (Extended Data Fig. 1d)^21^. In addition, we characterized the *in vivo NET1* mRNA distribution in stratified epithelial tissues of other species by examining primate tongue and human skin biopsies. Importantly, we again found in these cases that *NET1* mRNA exhibits a significant basal enrichment suggesting that basal *NET1* mRNA localization is evolutionarily conserved (Fig. 1a,d). We thus focused on the mouse *Net1* mRNA for further molecular and functional characterization utilizing the stratified tongue epithelium as an amenable experimental system.

The stratified squamous epithelium of the tongue consists of epithelial cells at different proliferation and differentiation states. Basal keratinocytes attached to the BM can enter the cell cycle and upregulate proliferating cell nuclear antigen (PCNA) during S/G2/M-phases^24^. To see whether *Net1* mRNA localization differs between quiescent and proliferating basal cells, we measured basal *Net1* accumulation in PCNA^Hi^ and PCNA^Lo^ cells (Fig. 1e). *Net1* distribution was not significantly different between cell proliferation states, based on this cell division marker (Fig. 1f). Basal Krt5+ cells can additionally transition to the suprabasal layer while upregulating differentiation markers like keratin 13 (Krt13) in the mouse tongue. As a result, such Krt13+ cells eventually lose contact with the BM. Quantification of *Net1* mRNA amount (by RNA-FISH) showed that *Net1* is downregulated in suprabasal differentiated keratinocytes (Extended Data Fig. 1e). Interestingly, we could detect cells that appear to be in the process of delaminating, since they exhibit basally-oriented Krt13 cellular extensions intercalating between Krt5+ cells (Fig. 1g). Such cells exhibited accumulation of *Net1* mRNA at the tips of these extensions, albeit infrequently (Fig. 1g inset). We interpret this to suggest that basal *Net1* mRNA localization is transiently maintained in delaminating keratinocytes concomitantly with an overall *Net1* downregulation during differentiation. Overall, these data suggest that *Net1* mRNA expression and localization is notably elevated in BM-contacting keratinocytes, regardless of their proliferation state. We therefore focused our attention on characterizing *Net1* mRNA and its function in basal keratinocytes.

### Net1 mRNA accumulates in protrusion-like structures that contact microfibrils at the dermal epidermal junction

Basal cells of the tongue reside at the interface between the oral epithelium and the underlying connective tissue. This interface in stratified epithelia has been primarily studied in the skin and referred to as the dermal-epidermal junction (DEJ). We will be using the same terminology here, given the parallels between oral and skin tissue^25^. The DEJ includes the basal plasma membrane and specialized adhesions called hemidesmosomes, which connect the epithelium to the BM and the dermal extracellular matrix (ECM) through fibrillar anchoring complexes^26,27^. The DEJ has important roles in maintaining the epithelium both by acting as a reservoir for signaling molecules and by providing structural support^28,29^.

We performed high-resolution confocal imaging of the DEJ, because of the pronounced basal accumulation of the *Net1* mRNA. We specifically visualized the basal cell membrane through staining with integrin a6 (Itga6), a core hemidesmosome component. Strikingly, high-resolution imaging revealed a high degree of topographical variability of the basal membrane which consisted of numerous micron- or submicron-wide protrusion-like structures that interdigitated with the BM (Fig. 2a; full serial optical slices of these structures can be viewed in Supplementary Videos 1 and 2). While these topographically variable regions have been observed incidentally in older literature^30,31^, the associated components or potential function of these structures is unknown^32^. Notably, the *Net1* mRNA localized within, and quite frequently at the tips of these structures (Fig. 2a; yellow arrowheads). Despite their smaller size, these keratinocyte structures appear morphologically similar to the *in vitro* cultured mesenchymal cell protrusions, where *Net1* and other protrusion-localized mRNAs have been previously studied^13–15^. These data thus raise the intriguing possibility that these uncharacterized keratinocyte structures reflect *in vivo* equivalents of *in vitro* protrusion-like structures and are analogously associated with localized mRNAs, including *Net1*.

**Fig. 2:**
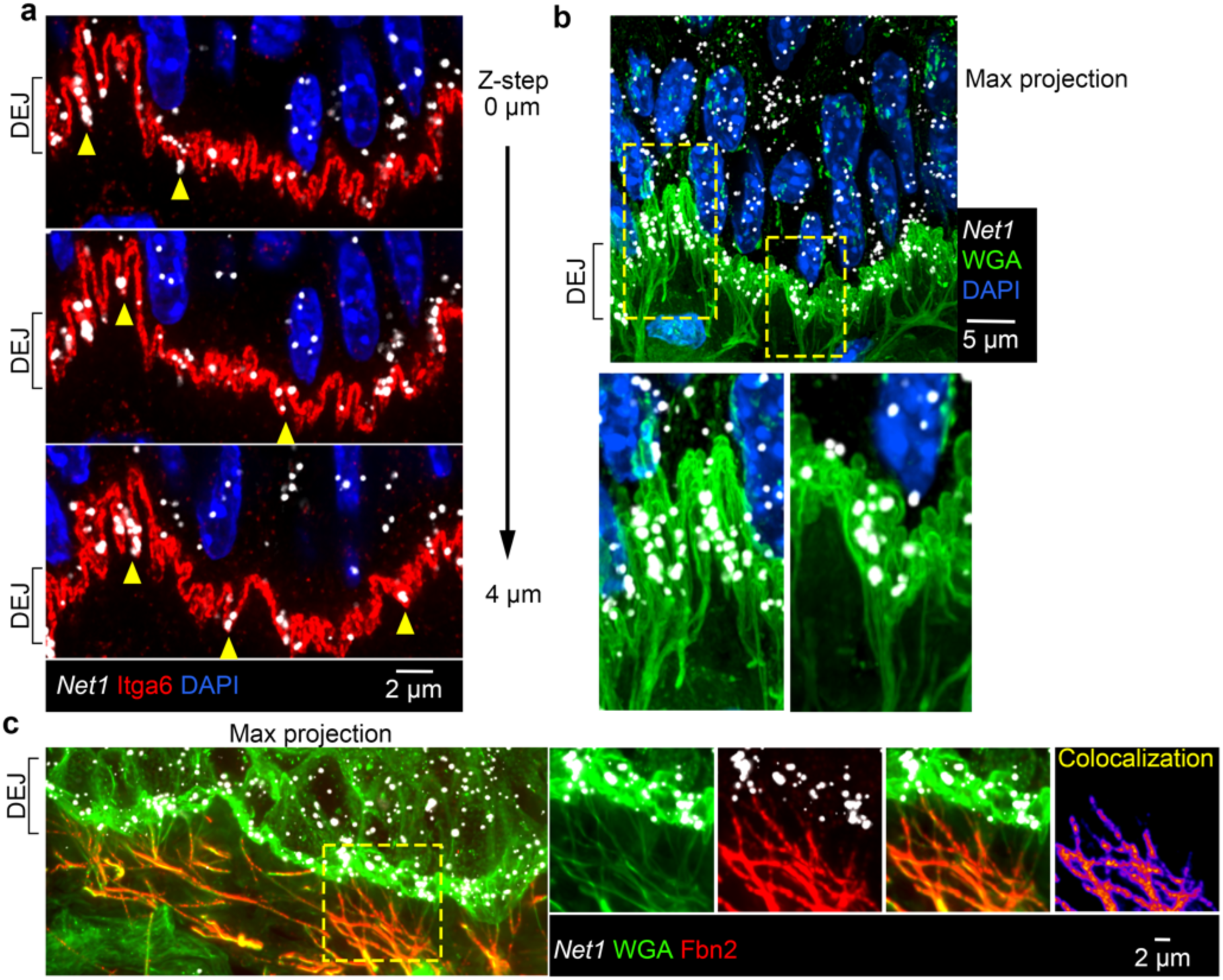
*Net1* accumulates in keratinocyte protrusion-like structures connected to microfibrils at the DEJ. **a**, Representative images of mouse tongue showing *Net1* at the DEJ. The basal keratinocyte cell membrane is visualized by Itga6 in sequential optical Z-slices. Arrowheads indicate *Net1* mRNA at protrusion-like structures. **b**, *Net1* and WGA staining of mouse tongue. Bottom panels: zoomed in regions, indicated by yellow boxes, highlighting fibers connecting with the basal keratinocyte cell membrane. Image is a max-intensity projection of ∼3 um in the z-axis. **c**, Mouse tongue section stained for *Net1* mRNA, WGA and Fbn2. Right panels show individual channels and overlay from boxed region. Colocalization panel shows pixel intensity overlap between WGA and Fbn2 channels.

Protrusion-like basal membrane structures can also be visualized upon staining with WGA (Fig. 2b; serial optical slices can be viewed in Supplementary Videos 1 and 2). Aside from visualizing the interdigitations this staining also revealed a complex fiber network that spans the dermis and appears to connect with the protrusion-like structures at the DEJ (Fig. 2b; note that this image presents a projection of multiple optical slices to allow better visualization of the fiber network). To determine what these WGA+ fibers correspond to, we surveyed various components of the DEJ and the dermal ECM. Co-detection with WGA indicated that these fibers are not enriched for hemidesmosome (Col17a1), basement membrane (laminin), or anchoring fibril (Col7) DEJ components (Extended Data Fig. 2). Because these fibers form an extensive network below the DEJ, we hypothesized that they are part of the dermal ECM. However, detection of collagen I (Col1a1), the primary dermal ECM component, showed that these fibers are not enriched for Col1a1 except at regions proximal to the DEJ (Extended Data Fig. 2). We discovered, though, that these fibers are strongly positive for fibrillin 2 (Fbn2), a major component of the microfibrils that are part of the elastic fiber network which allows transmission of mechanical forces across the dermis (Fig. 2c)^33–37^. Given the fact that stratified squamous epithelium is regularly mechanically deformed and that the microfibrils are essential in resisting this deformation, it is tempting to speculate that keratinocyte protrusions are sites of high mechanical stress. Pertinent to this, targeting of protrusion-localized mRNAs in *in vitro* cultures is coordinated with cellular mechanical state and influenced by ECM properties in various settings^11,16^. Therefore, the DEJ could reflect a physiological setting where the mechanoresponsive localization of mRNAs is particularly relevant.

### Net1 mRNA localization is necessary for maintaining epithelial architecture and protrusion-like structures at the DEJ

To broadly address the functional role of *Net1* mRNA localization in epithelial physiology, and protrusion-like structures specifically, we employed sequence blocking phosphorodiamidate morpholino oligos (PMOs). PMOs antisense to GA-rich regions of human protrusion-localized mRNAs have been shown to specifically prevent the localization of the targeted mRNA by blocking the formation of a transport-competent complex between the mRNA and the KIF1C kinesin^14,15,38^. To determine whether we can extend the use of this approach to mice, we designed PMOs that target the analogous GA-rich regions in the mouse *Net1* 3’UTR and tested them in *in vitro* cultured mouse NIH/3T3 fibroblasts. PMOs were either targeted to *Net1* GA-rich regions (PMOs #992 and #1016) or another downstream area (PMO #1620), or to an unrelated sequence (control PMO) (Extended Data Fig. 3a). Indeed, GA-targeting PMOs, but not other sequences, significantly prevented the localization of the *Net1* mRNA to the periphery of 3T3s without affecting another protrusion localized mRNA, *Cyb5r3* (Extended Data Fig. 3b, c). Importantly, and consistent with prior reports in human cells, the effect on *Net1* mRNA localization was not accompanied by any detectable change in the amount of *Net1* mRNA (Extended Data Fig. 3d)^15^. We also measured the enrichment of various mRNAs within isolated 3T3 protrusions and observed that Net1 PMO delivery affected solely the enrichment of the *Net1* mRNA while all other detected transcripts were unaffected (Extended Data Fig. 3e). We further validated that Net1 PMOs #992 and #1016 can similarly disrupt *Net1* mRNA localization in an immortalized keratinocyte cell line *in vitro* (Extended Data Fig. 4a,b), again without affecting *Net1* mRNA levels or the amount of Net1 protein produced (Extended Data Fig. 4c-e). These data confirm the broad conservation of the mRNA trafficking mechanisms to protrusions, and the applicability of PMO delivery as a specific tool to modify mRNA distributions in diverse cell types.

To assess the physiological importance of *Net1* mRNA localization *in vivo* we intradermally injected the two *Net1* localization altering PMOs or two control PMOs (one targeting GFP and one corresponding to a scrambled Net1 #992 sequence) into mouse tongues. PMOs were administered every 2 days over 6 days (Fig. 3a). We confirmed, by small RNA in situ hybridization, that PMOs reached and were taken up by the cells in the epithelium (Fig. 3b). Significantly, when we measured *Net1* mRNA distribution in injected tissues, there was a substantial reduction in the fraction of basal *Net1* mRNA upon administration of the GA-rich region targeting PMOs (Fig. 3c,d). Therefore, PMOs can prevent the basal accumulation of the endogenous *Net1* mRNA in epithelial tissues *in vivo*.

**Fig. 3:**
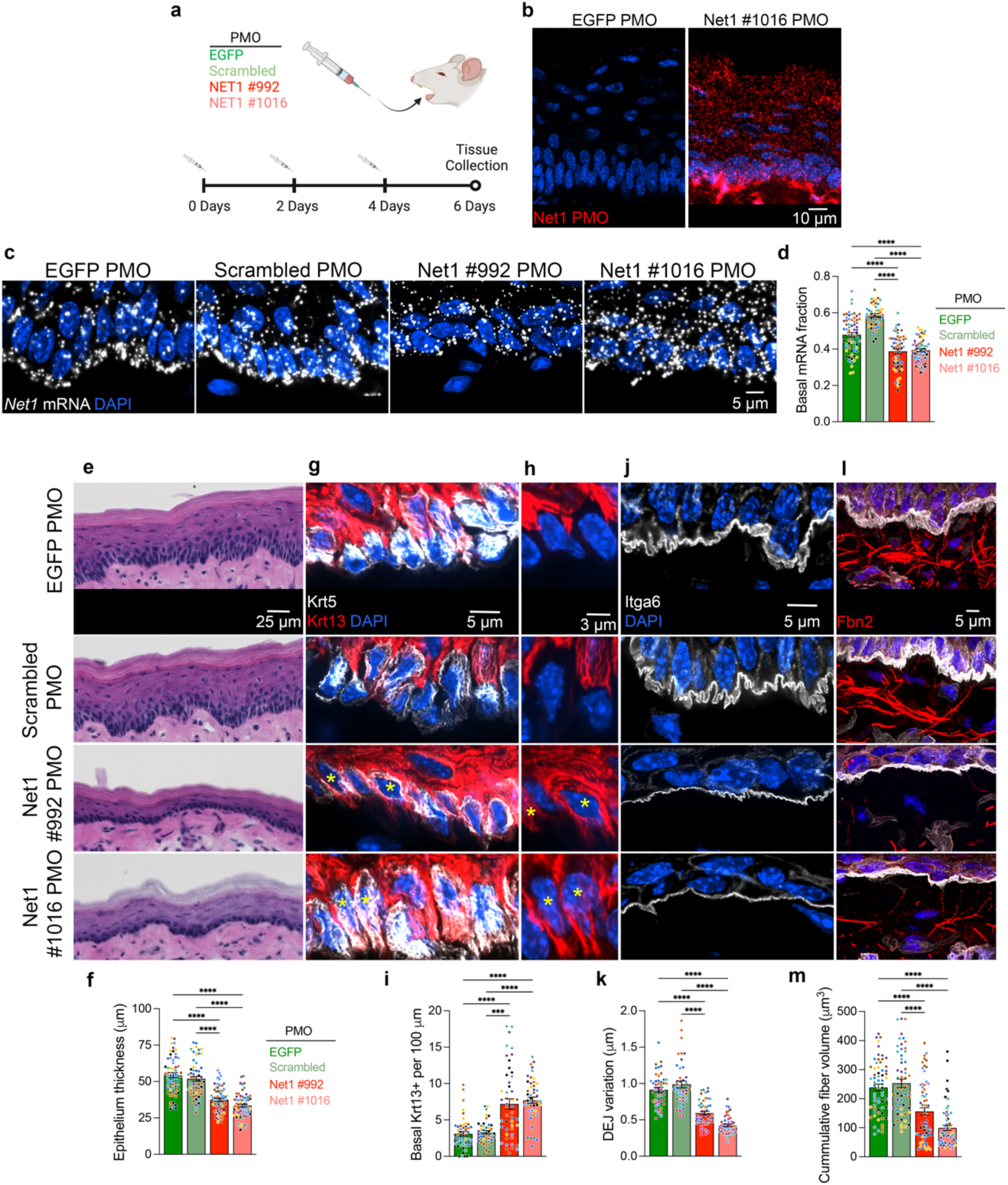
*Net1* mRNA localization controls epithelial homeostasis and keratinocyte interaction with the DEJ. **a**, Schematic of PMO injection protocol into the ventral side of mouse tongues using sequences targeting *EGFP* or the *Net1* 3’UTR at positions #992 and #1016, as well as a scrambled version of #992. **b**, PMO uptake into the tongue epithelium was visualized using Net1 #1016 PMO-specific probes. **c**, Visualization of *Net1* distribution in PMO injected tongues. **d**, Basal *Net1* mRNA fraction in PMO treated tissues. **e**, Visualization of overall oral epithelium architecture using H&E on ventral portion of the tongue. **f**, Measurement of tongue epithelial thickness. **g**, Visualization of basal (Krt5) and spinous (Krt13) keratinocytes in the basal compartment of tongue epithelium. **h**, Select areas from (g). Krt13 channel is only shown to highlight examples of Krt5+/Krt13-basal cells in control PMO treated tongues and Krt5+/Krt13+ in Net1 PMO treated tongues. Asterisks indicate double positive cells. **i**, Number of Krt5+/Krt13+ cells in the basal compartment normalized to DEJ length. **j**, Visualization of DEJ using Itga6 in PMO treated mouse tongues. **k**, Measurement of DEJ topographical variation. **l**, Visualization of Fbn2+ fibers in PMO-treated tongues. **m**, Quantification of DEJ proximal Fbn2+ fiber volume. Bar graph data are measurements from individual tissue regions (ROIs) with mean ± SEM; n = 42-64 ROIs from N = 7-8 mice per condition. Data from individual mice are represented with different colors. *** P ≤ 0.001 and **** P ≤ 0.0001 by Brown-Forsythe and Welch ANOVA followed by Dunnett’s T3 multiple comparison test (d) or Kruskal-Wallis ANOVA followed by Dunn’s multiple comparison tests (f,i,k,m).

Further phenotypic analysis revealed prominent defects in epithelial architecture detected by H&E staining (Fig. 3e). Specifically, the epithelium of Net1 PMO-treated tongue epithelia was thinner in comparison to control treated tissues (Fig. 3f). Such changes in epithelial architecture can be brought about by defects in cellular differentiation^39,40^. Visualization of canonical markers of the basal and spinous compartment, Krt5 and Krt13 respectively, revealed that Net1 PMO-treated tongue contained significantly more Krt5/Krt13+ double positive cells in the basal layer (Fig. 3g-i). This suggests either spontaneous differentiation of Krt5+ cells or alternatively a defect in cell detachment from the basement membrane.

Given the fact that *Net1* mRNA localizes to protrusion-like structures at the DEJ, we investigated whether altering *Net1* localization from the basal surface affects DEJ organization. Indeed, Net1 PMO-treated epithelium exhibited drastic flattening of protrusion-like structures formed by basal keratinocytes (Fig. 3j). We quantified this change by measuring topographical variation of the DEJ using optical slices of the Itga6-stained basal membrane (Extended Data Fig. 5). Using this metric, DEJ variation was halved in Net1 PMO-treated epithelia (Fig. 3k). As shown above, basal protrusions appear to be connected to Fbn2+ fibers. We thus asked whether altering *Net1* localization has broader effects on the microfibrils near the DEJ. Consistent with this, the volume of Fbn2+ fibers was significantly reduced in regions proximal to the DEJ (Fig. 3l,m). This result could reflect ECM remodeling near the epithelium but may also be due to decreased antibody-epitope accessibility on individual fibers due to altered mechanical strain, which can lead to changes in fibrillin domain folding^41^. Regardless, it is notable that dysregulation of the DEJ during aging or in genetic diseases, including ones caused by mutations in fibrillins, has strong negative implications on epidermal physiology^32,33,37,42^. Altogether, these results demonstrate that altering the basal localization of a single mRNA, *Net1,* is sufficient to drastically influence keratinocyte physiology and tissue homeostasis, potentially by altering interactions of the basal epithelium with the DEJ.

### Net1 mRNA localization controls Net1 protein distribution and activity in vivo

To understand the mechanism through which altering *Net1* mRNA localization affects keratinocyte physiology, we considered that *Net1* mRNA location might affect the activity of the encoded Net1 protein as a RhoA GEF. Net1 is quite unique among RhoA regulators in that it is controlled by nucleo-cytoplasmic trafficking^43,44^. While RhoA activation largely occurs at the plasma membrane, Net1 can be imported and sequestered in the nucleus, providing an “off-switch” for Net1-dependent RhoA activation^15,18^. *Net1* mRNA location balances Net1 nuclear import versus cytoplasmic retention by a partner-selection mechanism in mesenchymal cells *in vitro*. Specifically, translation of Net1 from protrusion-localized mRNA favors its interaction with a membrane scaffold and promotes RhoA activation by Net1^15^. To determine whether this mRNA location-dependent regulation operates in the tongue epithelium *in vivo,* we looked at the subcellular distribution of the Net1 protein. We first validated the specificity of the Net1 antibody, by immunofluorescence staining of immortalized keratinocytes upon Net1 knock down with two different siRNAs (Extended Data Fig. 6a,b). We then observed Net1 protein distribution in tongue sections. Net1 protein signal was primarily detected in the basal epithelial layer (Fig. 4a), consistent with the basal cell specific expression pattern of *Net1* mRNA, mentioned above (Extended Data Fig. 1e). This further supports the specificity of Net1 protein detection. We note that while Net1 protein is primarily nuclear in cultured keratinocytes, in *in vivo* tissue Net1 is primarily cortical (compare Fig. 4a and Extended Data Fig. 6a). While the basis for this differential regulation is still unclear, this has been also observed in other studies^18,45,46^. Additionally, Net1 is observed throughout the basal keratinocyte cortex (Fig. 4a), consistent with *in vitro* results showing that Net1 retained in the cytoplasm can distribute away from the site of protein synthesis^15^. To quantify changes in Net1 protein distribution we measured its cortical/non-cortical intensity (Fig. 4a-c). Net1 PMO-treated epithelium showed a reduced cortical Net1 accumulation (Fig. 4d), indicating that *Net1* mRNA location controls Net1 protein distribution *in vivo*.

**Fig. 4:**
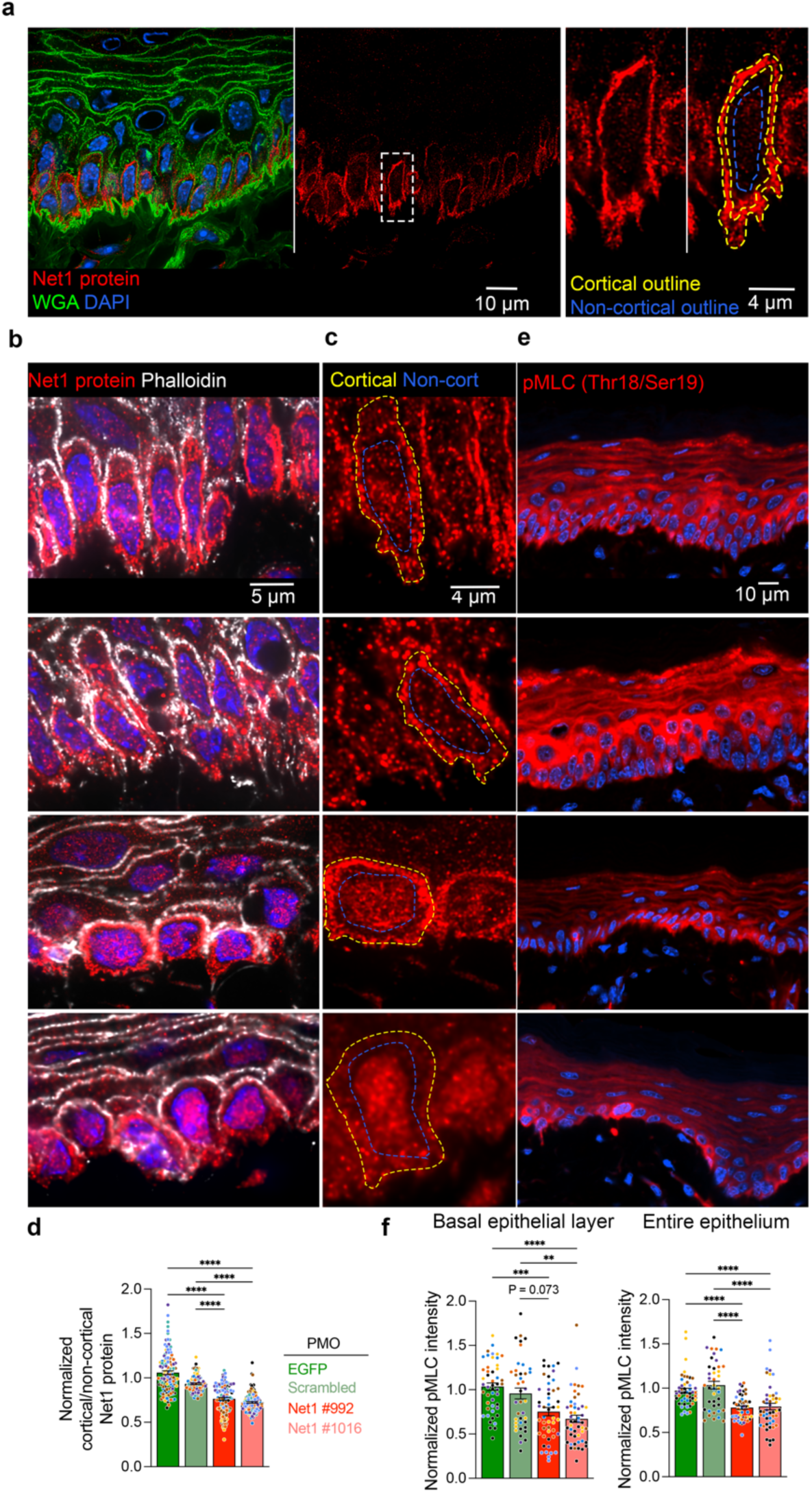
*Net1* mRNA localization regulates Net1 protein distribution and activity *in vivo*. **a**, Visualization of Net1 protein and WGA in mouse tongue keratinocytes. Left panel: overlay. Middle panel: Net1 channel showing predominant expression in basal keratinocytes. Right panels: zoom in of boxed region. Dashed yellow and blue lines indicate cortical and non-cortical regions, respectively, used for Net1 distribution measurements. **b**, Visualization of Net1 protein with phalloidin as a marker of the cell cortex in PMO treated tissues. **c**, High magnification visualization of Net1 protein where the cell cortex and non-cortical region are indicated by yellow and blue broken lines, respectively. **d**, Quantification of Net1 protein distribution (ratio of cortical to non-cortical mean intensity). **e**, Visualization of MLC2 phosphorylation (pMLC) in the mouse epithelium. **f**, Quantification of mean pMLC2 fluorescent intensity for basal keratinocytes (left) and the entire oral epithelium (right). Bar graph data are individual measurements with mean ± SEM; n=105-120 cells (d) or 40-48 ROIs (f) from N = 7-8 mice per condition. Data from individual mice are represented with different colors. ** P ≤ 0.01, *** P ≤ 0.001, **** P ≤ 0.0001 by Kruskal-Wallis ANOVA followed by Dunn’s multiple comparison test.

A reduction in cortical Net1 amount would be predicted to result in reduced activation of the RhoA GTPase. Given that it is technically difficult to directly measure RhoA activity in tissues, we examined the phosphorylation of myosin light chain (pMLC), a major downstream RhoA target^47^. pMLC levels were visualized using a phospho-specific antibody under conditions that allow specific detection (Extended Data Fig. 7). Quantification of pMLC levels in Net1 PMO-injected tongue epithelium revealed that altering *Net1* mRNA localization reduced pMLC in both basal keratinocytes and more broadly across the entire epithelium (Fig. 4e,f). Therefore, preventing the basal localization of the *Net1* mRNA leads to altered Net1 protein distribution and a reduction in RhoA-pMLC signaling. Given that this pathway is well known to control cytoskeletal tension and cell attachment, it is likely that it reflects the mechanism linking *Net1* mRNA targeting to DEJ structure and epithelial organization.

### DEJ morphology is sensitive to fluctuations in basal keratinocyte cytoskeletal state

To test this idea, we tried to independently address whether changes in RhoA signaling within basal keratinocytes control their interaction with the DEJ. For this, we used a genetic system for cell-type specific RhoA activation^48,49^. Krt14-rtTA mice were crossed with a genetic knock-in of a constitutively active RhoA GEF, ArhGEF11 (ArhGEF11^CA^), under the control of a tetracycline-inducible promoter (Fig. 5a), to achieve RhoA activation specifically in basal keratinocytes. Doxycycline injection led to basal keratinocyte-specific overexpression of ArhGEF11^CA^ in mouse tongue, visualized by an HA-tag, as early as 10hrs post-injection (Fig. 5b). ArhGEF11^CA^-expressing cells exhibited increased cortical pMLC and filamentous actin (phalloidin staining) indicative of RhoA-induced cytoskeletal tension (Fig. 5b).

**Fig. 5:**
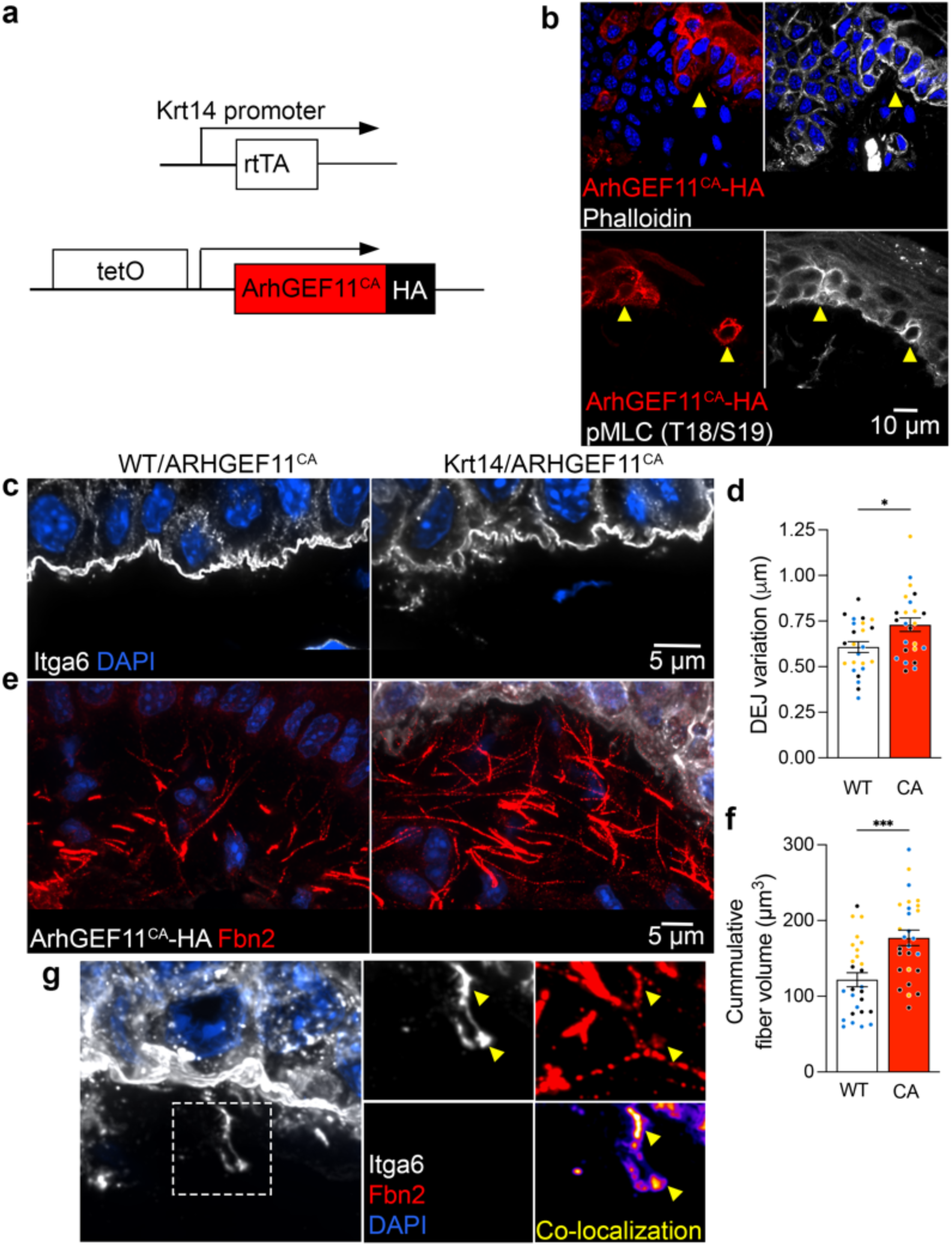
RhoA activation in basal keratinocytes remodels the DEJ. **a**, Schematic of ArhGEF11^CA^-HA expression under the control of tetracycline-inducible promoter where the reverse tetracycline-controlled transactivator is expressed in basal epithelial cells (Krt14-rtTA). **b**, Visualization of the actin cytoskeleton (phalloidin) and pMLC in ArhGEF11^CA^-HA-expressing cells. Mosaic expression of ArhGEF11^CA^ is indicated by yellow arrowheads. **c**, Visualization of the DEJ by Itga6 in WT or ArhGEF11^CA^ mice. **d**, Quantification of DEJ variation in knock-in mice. Measurements in ArhGEF11^CA^ (CA) mice were made in regions with continuous HA expression in basal keratinocytes across an analysis ROI. **e**, Visualization of HA and Fbn2 at the DEJ of WT or ArhGEF11^CA^ mice. **f**, Quantification of Fbn2 volume proximal to the DEJ. **g**, Left panel: example of an ArhGEF11^CA^-expressing cell exhibiting extension of the plasma membrane below the DEJ after supraphysiologic induction of RhoA activation. Plasma membrane is stained through Itga6. Right panels: Boxed region showing also Fbn2 staining and co-localization between the two channels. Yellow arrowheads indicate areas of high Itga6-Fbn2 co-localization. Bar graph data are measurements from individual ROIs with mean ± SEM; n = 24 (d) or 27 (f) ROIs from N = 3 mice. Data from individual mice are represented with different colors. * P < 0.05, *** P < 0.001 by unpaired two-tailed Student’s *t*-test.

We examined tongue tissue architecture in these mice, comparing them to mice with the ArhGEF11^CA^ cassette that were not crossed with Krt14-rtTA mice (WT/ArhGEF11^CA^). We focused our quantitative analysis on DEJ regions where all of the basal keratinocytes express the HA-tagged ArhGEF11^CA^. Remarkably, this short-term induction of RhoA signaling was sufficient to significantly increase the topographical variation of the basal plasma membrane at the DEJ (Fig. 5c,d). Concomitantly, DEJ proximal Fbn2+ fibers became more prominent and displayed a higher overall volume in regions proximal to the epithelium (Fig. 5e,f). Given the short induction time we favor the idea that increased fibrillin detection reflects, at least partly, increased epitope accessibility through tension-induced unfolding, rather than solely an increase in their absolute amount.

A unique feature observed upon ArhGEF11^CA^ expression was the occasional appearance of Itga6 strands co-localizing with Fbn2+ fibers far below the overall level of the basal plasma membrane (Fig. 5g). The fact that these do not appear under normal conditions suggests that they are formed upon the supraphysiologic tension induced under our experimental setting. We envision that fibrillin resistance to keratinocytes’ tensile forces eventually leads to fiber retraction away from the epithelium. In this scenario the residual tension in fibrillin fibers pulls portions of the plasma membrane below the ordinary DEJ (Fig. 5g). The tight colocalization of Itga6 with fibrillin suggests the existence of stable mechanical coupling between the keratinocyte plasma membrane and microfibrils. These data provide orthogonal evidence that changes in RhoA activity, and cytoskeletal tension, is sufficient to alter keratinocyte connections with the DEJ. Therefore, the RhoA pathway is the likely mediator of the ultrastructural changes at the DEJ in response to altering *Net1* mRNA localization. Overall, the evidence presented here reveals mRNA localization as an unappreciated level of post-transcriptional regulation that controls the mechanical coupling of the epithelium with the underlying connective tissue, which broadly affects tissue physiology.

## Discussion

Post-transcriptional mRNA regulation has well known roles both in normal physiology as well as in disease, exemplified for instance by miRNA-mediated regulation during development^50^, or the therapeutic modulation of splicing patterns^51^. Targeting of mRNAs to specific subcellular domains has emerged as another widespread level of post-transcriptional mRNA control with the potential to spatially alter the proteome at a subcellular level^3,4^ and control protein function^15^. Nevertheless, the contribution of mRNA localization to mammalian physiology has remained largely unexplored. Here, we demonstrate that the subcellular localization of an individual mRNA, *Net1*, is an important regulator of epithelial tissue homeostasis. This observation underscores the importance of this post-transcriptional regulatory mechanism in epithelial tissue physiology where ∼15-30% of the transcriptome is differentially polarized^1,2^ and sensitive to physiological stimuli^1^. Notably, we show that the basally localized *Net1* mRNA is associated with protrusion-like structures of keratinocytes that contact a mechanosensitive network of dermal microfibrils. We speculate that these underappreciated structures at the DEJ may be mechanical connections necessary for long range force transmission between the epidermis and dermis. Targeting of the *Net1* mRNA at these DEJ structures is necessary for their formation and its loss results in remodeling of the dermal ECM. Our evidence further suggests that these effects are mediated through basal cell autonomous activation of RhoA. Indeed, Net1-mediated activation of RhoA is necessary for collagen I remodeling in breast tissue^52^ as well as basement membrane breakdown during embryonic development^45^. The findings presented here indicate that such RhoA-regulated cell-ECM interactions are mediated at least partly by localized translation of the *Net1* mRNA. Interestingly, *Net1* mRNA localization is itself mechanosensitive, being affected by the properties of the ECM^11,16^, implying the existence of feedback regulation that maintains an RNA-dependent mechanical homeostasis of the DEJ. This maintenance of homeostasis extends to keratinocyte function, including differentiation, which is regulated by cell-autonomous and tissue-wide variation in mechanical state^39,40^.

Alterations in DEJ components, including breakdown of the microfibrillar network and a morphological attenuation of keratinocyte protrusions, occurs during aging^32,42^. Genetic conditions, termed fibrillinopathies, also affect fibrillins and have deleterious effects on epidermal homeostasis^29,37,53^. We speculate that changes in basal localization of *Net1*, and likely other protrusion mRNAs, might be a contributing component in the mechanical uncoupling of the epidermis in aging and disease. Given that Net1 is widely expressed and consistently localized in a variety of tissues it is possible that its unique regulatory mechanism is utilized for maintenance of epithelium-ECM connections in additional tissues. Furter understanding of the dynamic control and functional contributions of such events occurring at tissue interfaces will provide exciting new directions and context for studying RNA transport and local regulation in mammalian physiology.

## Acknowledgments

We thank the CCR Genomics Core of the National Cancer Institute, NIH for ddPCR and nanoString nCounter analysis, and the microscopy facility of the Laboratory of Cancer Biology and Genetics for Axioscan imaging. This research was made possible in part using biomaterials from the NIA Nonhuman Primate Tissue Bank (https://www.nia.nih.gov/research/dab/nonhuman-primate-tissue-bank) at the Wisconsin National Primate Research Center, University of Wisconsin-Madison under contractual agreement with the National Institute on Aging (NIA). This work was funded by the Intramural Research Program of the Center for Cancer Research, National Cancer Institute (NCI), National Institutes of Health (NIH) (1ZIA BC011501 to S.M., ZIA BC 011763 to R.B., and ZIA BC 011682 to R.W.), and by R01-AR067203 and R01-AR081081 to T.L..

## Author Contributions

D.E.M. and S.M. conceived the project and designed experiments. D.E.M. and T.D.M. performed experiments. D.E.M., A.N.G., N.J., T.L., R.W., R.I-B. and S.M. analyzed data. S.M. and D.E.M. wrote original manuscript. All authors reviewed and edited the manuscript.

## Declaration of Interests

The authors declare no competing interests.

## Data and Code availability

Previously published sequencing data that were reanalyzed here are available under accession codes (GSE129218 and GSE77197). All other data supporting the findings of this study and custom analysis codes are available from the corresponding authors on reasonable request.

## Methods

### Animal handling and tissue collection

All experiments involving animals were performed in accordance with guidelines set by the Institutional Animal Care and Use Committees of the National Cancer Institute (National Institutes of Health, Bethesda, MD) and Duke University (Durham, NC; Protocol #: A255-23-12). All mice (4-12 weeks) were housed in sterile filter-capped cages, fed and watered ad libitum, in a 12:12 light-dark cycle animal facility.

For injections, custom octaguanidine dendrimer conjugated PMOs (Vivo-Morpholino, GeneTools) that target *EGFP* or *Net1* mRNA were synthesized (Extended Data Table 1). PMOs (5 nmol/inj) and fluorescently labelled dextran (2.5 μg/inj.; Molecular probes; D1976) were mixed in isotonic sterile saline. 10µL of the solution was subcutaneously injected in the ventral portion of the tongues of anesthetized male and female C57BL/6/NCr mice (Charles River; age 10-12 weeks). Anesthesia was performed by brief exposure to 3% isoflurane followed by intraperitoneal injection of 100mg/kg ketamine and 10 mg/kg xylazine. The tongues were injected with PMOs every two days for a total of three injections. Mice were sacrificed for tissue collection 6 days after the initial injection.

4-week-old TRE-Arhgef11/Krt14-rtTA mice (3 males and 3 females) were IP-injected with 25 mg/kg doxycycline (Sigma) 10 hours before sacrifice and tissue collection.

Tissue was collected from male and female euthanized mice (age 4-12 weeks) washed gently in PBS then flash frozen in optimal cutting temperature compound (OCT; Fisher Healthcare; 4585). Tissue embedded OCT blocks were cut 8-10um thick on a Leica cryostat microtome CM1860. Tissues were collected on positively charged slides (Rankin; 20290W) then stored at - 80C. The gross tissue morphology of each tissue was visualized by H&E staining (abcam; ab245880).

### Mouse keratinocyte cell line generation

Mouse keratinocytes were isolated from tail skin of mixed background mice (FVB/N and C57) by physical separation of the epidermis and dermis from the hypodermis. Tail skin was sterilized with 10% v/v betadine/iodine in PBS. Excised tissue was digested overnight at 4C using 2U/mL of dispase II (StemCell; 07913) diluted in .25% trypsin without EDTA (Gibco; 15050065). After digestion the tissue was minced and mixed with media and antibiotics. Keratinocytes were then isolated from the digest by sequential straining using 100 μm and 40 μm cell strainers. Cells were cultured for one passage using EpiLife medium with 60 μM calcium (Gibco, Waltham, MA, MEPI500CA), supplemented with Human Keratinocyte Growth Supplement (HKGS, Life Technologies, S0015), mouse EGF (10 ng/ml, R&D Systems, Minneapolis, MN, 2028EG200), and Y-27632 compound (10 μM, Tocris Bioscience, Bristol, United Kingdom, 12-541-0). Keratinocytes were then immortalized by infecting with lentiviruses expressing SV40 Large T Antigen (Addgene Plasmid #170255). Cells were selected with Hygromycin (200 ug/mL) for two days and then cells were passed for two passages in EpiLife and described supplements. Then cells were cultured in DMEM (Sigma-Aldrich Inc) containing 10% fetal bovine serum (FBS) (Sigma-Aldrich Inc), antibiotic/antimycotic solution (Sigma-Aldrich Inc), mouse EGF (10 ng/ml, R&D Systems, Minneapolis, MN, 2028EG200), and Y-27632 compound (10 μM, Tocris Bioscience, Bristol, United Kingdom, 12-541-0), for at least two passages before experiments. Lentiviruses were produced by transfecting Lenti-X 293T cells with pMD2.VSVG (Addgene plasmid #12259) and psPAX2 (Addgene plasmid #12260). Lenti-X 293T cells were obtained from Takara Bio and cultured in DMEM (Sigma-Aldrich) containing 10% fetal bovine serum (Sigma-Aldrich).

### Immunofluorescence and tissue staining

Fresh frozen sections or cultured cells were fixed in 4% formaldehyde (Sigma; F8775) for 15-30 minutes at RT washed 3×2 minutes in PBS and permeabilized in 0.2% Triton-X-100 for 5-20 minutes. Sections were washed in PBST and blocked for 1 hour in 3% bovine serum albumin (BSA) and 2% goat serum (GS) in PBS. Antibodies were diluted in blocking buffer and incubated overnight (Extended Data Table 2). Samples were washed 3×2minutes in PBST and incubated with fluorophore conjugated secondary antibodies corresponding to the host species (Extended Data Table 2; 1:500 in blocking buffer). Samples were washed with PBST 3×2 minutes then counterstained with a fluorophore conjugated variation of wheat germ agglutin (Invitrogen; W11261; 10ug/mL), phalloidin (Invitrogen; A12379), and/or DAPI. Tissue samples were mounted using #1.5 coverglass (Fisher; 12544D) with either prolong glass or diamond mounting medium (Invitrogen; P36980 or P36962) and allowed to set overnight at room temperature.

Phosphorylated-myosin light chain 2 (Thr18/Ser19) was detected in tissue samples using a modified immunofluorescence protocol. Briefly samples were fixed in 4% formaldehyde for 3 hours at RT, washed 3×5 minutes in TBS then treated with 1% SDS diluted in TBS for 20 minutes at RT with regular agitation. Samples were washed thoroughly in 3×5 minutes in TBST (0.1% Tween-20). Samples were blocked in 3%BSA and 2% GS in TBS for 1 hour. pMLC specific rabbit mAb (Cell signaling; 95777) was diluted in blocking buffer and incubated at 4C overnight. Staining was completed as described above with TBS instead of PBS.

Specificity of the pMLC antibody was validated by pre-treating tissues with 4000U λ phosphatase (NEB; P0753S) diluted in 1xNEBuffer for protein phosphatases and 1mM MnCl_2_ overnight at 30℃. Samples were washed thoroughly in TBST before proceeding with the staining protocol described above.

### Cell culture

NIH/3T3 mouse fibroblast cells (ATCC) were grown in Dulbeco’s Modified Eagle Medium (DMEM; Gibco; 11995-065) supplemented with 10% calf serum (Cytiva; SH30087.04), sodium pyruvate, and penicillin/streptomycin (Gibco; 15140122). Immortalized mouse keratinocytes were cultured in DMEM supplemented with 10% Fetal Bovine Serum,10ng/mL mouse EGF, and 10 μM Y27632. Cells were cultured at 37°C, 5% CO_2_ and passaged with either 0.05% (3T3) or 0.25% (keratinocytes) trypsin with EDTA (Gibco; 25300054 or 25200056). Cells used in this study have tested negative for mycoplasma.

For PMO delivery, cells were grown to 70% confluence in 12 well plates then transfected with the indicated PMOs at a concentration of 15 μM in basal media. For small interfering RNA (siRNA) experiments cells were cultured to 50-70% confluency in 6 or 12 well plates then transitioned to antibiotic free media. Cells where transfected with the indicated siRNA (Qiagen; final concentration 40 nM; Extended Data Table 1) which were pre-complexed with Lipofectamine RNAimax (Invitrogen; 13778075) transfection reagent according to the manufacturer’s instructions. Cells were assayed for protein expression 48 hours after transfection.

### Imaging

Fixed samples were imaged on a Nikon Eclipse Ti2-E inverted microscope with a Yokogawa CSU-X1 spinning disk confocal scanner unit and operated using NIS-Elements software. Images were acquired using 20x (Plan Apo 20x; NA = 0.75; WD = 1000μm), 40x (Plan Fluor 40; Oil; NA = 1.30; WD = 240 μm), 60x (Apo 60x λS DIC N2; Oil; NA = 1.40; WD = 140 μm), or 100x (Apo TIRF DIC N2; Oil; NA = 1.49; WD = 120 μm) objectives and a Hamamatsu ORCA-Fusion BT Gen III back-illuminated sCMOS cameras. Images were denoised and deconvolved using the Richardson-Lucy algorithm in the NIS-Elements analysis software.

Whole sample brightfield images of H&E-stained samples were taken using a Zeiss Axio Scan.Z1 (Zeiss) equipped with a Hamamatsu OrcaFlash 4.0 camera using 10x and 20x objectives. Images were aquired using the Carl Zeiss Zen 2.3 software and analyzed using the HALO image analysis platform (Indica Labs).

### Basal mRNA quantification

mRNA accumulation to the basal region of epithelial layers was quantified by manually segmenting a maximum projection of ∼1 μm thick optical sections in MATLAB (Mathworks; R2021b-R2023b). A custom script was used to separate the segmented layer into 10 evenly divided bins from the basal to apical side of a layer, and to measure fluorescent intensity in each bin after a uniform background subtraction. The fraction of intensity in each bin was then calculated per layer and used to measure relative fraction of basal mRNA (bottom 30% of a segmented layer).

### Net1 protein distribution

Net1 protein distribution was calculated on a per cell basis by manually segmenting basal keratinocytes in the mouse tongue using the FIJI image analysis platform^54^. Segmentation was performed on individual optical slices taken with a 100x objective described above. Cortical Net1 protein mean intensity was measured in the peripheral 15% of the cell then normalized to the most central 65% of the segmented cell.

### DEJ variation analysis

To measure the complex protrusion-like structures at the DEJ we decided to use a metric that incorporates DEJ topographical variation (the full process is visualized in Extended Data Fig. 5). Briefly, the DEJ was manually segmented into individual optical slices using the FIJI line tool (120 pixel centered on the DEJ). Itga6 fluorescent intensity was automatically thresholded and skeletonized. The basal cell membrane was estimated for each x-position using the skeletonized image. The x- and y- coordinates from these measurements were then used to create a 2D plot profile from which the absolute distance between the basal cell membrane and the segmentation line was measured for each x-coordinate. Distances were summed and normalized to the number of x-positions measured to yield a single DEJ variation metric. This analysis approach is analogous to the surface characterization strategies common in machining applications^55^.

### Fibrillin 2+ quantification analysis

Fibrillin 2+ fiber volume was measured using 4 μm thick optical stacks acquired from epidermis proximal dermal areas (∼50 μm^2^) using a 100x objective. Imaris (Bitplane; v9.9.0) was used to generate Fbn2+ fiber volumes by thresholding Fbn2 fluorescence intensity equally between samples within a given experiment. Morphological filters were then equally applied across conditions to remove small spherical volumes that were not consistent with fiber segments. The cumulative fiber volume was then measured from all extracellular Fbn2+ fiber volumes in each image.

### Peripheral distribution index analysis

FISH images of *in vitro* cultured cells were analyzed using a previously published MATLAB script^56^. Briefly, the peripheral distribution index (PDI) of individual RNAs was measured by calculating the difference between the geometric centroid of a cell and the intensity weighted centroid of an RNA. This value is then normalized to a hypothetical uniformly distributed RNA. This ratio represents the relative peripheral distribution of an RNA in the context of a cell’s geometry. RNAs are considered peripheral if PDI values are greater than 1, homogenous if equal to 1, and perinuclear if less than 1.

### RNA detection

Detection of RNA in tissues sections was performed using the RNAscope Multiplex Fluorescent Reagent kit v2 (ACD; 323100). Tissues were first pre-processed as follows: fresh frozen tissues were fixed for 45 minutes in 4% formaldehyde (Sigma) diluted in PBS at room temperature. Samples were washed 3×2 minutes in PBS then sequentially dehydrated in EtOH (50, 70, 100%) and rehydrated in hydrogen peroxide (ACD). Samples were washed twice in distilled water then PBST (0.1% Tween-20). If proteins were co-detected, then antibodies were diluted in Co-Detection Antibody Diluent (ACD; 323160). Antibodies were incubated overnight at 4C after the hydrogen peroxide step. Samples were washed 3×2 minutes in PBST. Then the antibodies were fixed for 30 minutes using 4% formaldehyde then washed 3×2 minutes in PBST. Protease IV (ACD) was applied for 30 minutes at RT washed 3x in deionized water. Primary RNA probes (Extended Data Table 3) were applied, and the detection protocol was completed according to manufacturer’s instructions. Tissues were then counterstained with AF488 conjugated wheat germ agglutin, fluorescently labelled secondary antibodies, and DAPI.

PMOs were detected using the RNAscope Plus smRNA-RNA HD detection kit (ACD; 322780) according to the protocol described above and the manufacturer’s instructions. PMO detection probes were used at 1:1,000 of the manufacturer’s recommended concentration.

For RNA detection in NIH/3T3 cells or mouse keratinocytes, cells were plated on Collagen IV-coated (10 μg/mL; Sigma; C5533) #1.5 glass coverslips for 2-3 hours at 37C. Cells were fixed for 20 minutes at RT in 4% paraformaldehyde (EMS; 15710) then processed using the ViewRNA ISH Cell assay kit (Thermo Fisher Scientific; QVC0001) according to the manufacturer’s instructions and probes for individual mRNAs (Extended Data Table 3). Cell mask and DAPI were used as counterstain to detect the cell periphery and nucleus. Samples were mounted in prolong gold.

### RNA isolation and ddPCR

Bulk RNA was isolated from *in vitro* cultured cells using either Trizol LS (Thermo Fisher Scientific; 10296010) or RNeasy Plus Mini kit (Qiagen; 74134). 1μg of RNA was reverse transcribed using the iScript cDNA Synthesis Kit (Bio-Rad; 1768891) and used for droplet digital polymerase chain reaction (ddPCR). PCR reactions were prepared with cDNA, gene specific primers, and ddPCR EvaGreen Supermix (Bio-Rad; 186-4034). Droplets were generated from this reaction using the Automated Droplet Generator (Bio-Rad; 186-4101), PCR was then performed on a C1000 Touch Thermal Cycler (Bio-Rad; 185-1197), and droplet reading was performed on QX-200 Droplet Reader (Bio-rad; 186-4003). Results were quantified using the QuantaSoft software (Bio-rad).

### Protrusion/cell body isolation

Protrusion and cell bodies were isolated from serum-starved NIH/3T3 cells plated on transwell inserts with 3.0 μm porous polycarabonate membrane (Corning) as previously described^11^. Extracts were isolated using Trizol LS. Isolated RNAs were then quantified using the nanoString nCounter analysis platform and a custom-made codeset, according to the manufacturer’s instructions.

### Western blot

For western blotting the following primary antibodies were used: rabbit anti-NET1(1:1000; Bethyl; A303-138A) and mouse anti-⍺-tubulin (1:2000; Sigma; T6199). Anti-rabbit and anti-mouse secondary antibodies from Li-Cor were used at 1:10,000. Membranes were scanned using an Odyssey fluorescent scanner (Li-Cor) and bands were quantified using ImageStudioLite (Li-Cor).

### Human and primate tissues

De-identified human samples were collected from normal tissue following Mohs surgery for skin cancer removal. Frozen skin samples were collected and frozen in OCT before sectioning and confirmed to be normal by H&E staining. Under National Institutes of Health protocols, the use of biospecimens from de-identified discarded human tissue does not meet the regulatory criteria for human subject research and therefore institutional review board review or informed consent are waived.

Non-human primate tissues were obtained from the National Institute on Aging’s (NIA) Nonhuman Primate Tissue Bank. Fresh frozen tongue biopsies were obtained from 14–24 year-old female and male *Papio Anubis* (Olive Baboons) in the process of necropsy.

### Statistical analysis

Where relevant, data are shown as mean ± standard error of the mean (SEM). Individual measurements are shown with differing colors corresponding to individual experiments or animals. For statistical comparisons parametric tests were used to evaluate significant differences in normal distributed, homoscedastic datasets. Where either of these assumptions were not true, we instead performed non-parametric tests suitable for the data distribution. Specific statistical tests for each experiment are described in the figure captions.

---

**Extended Data Fig. 1:**
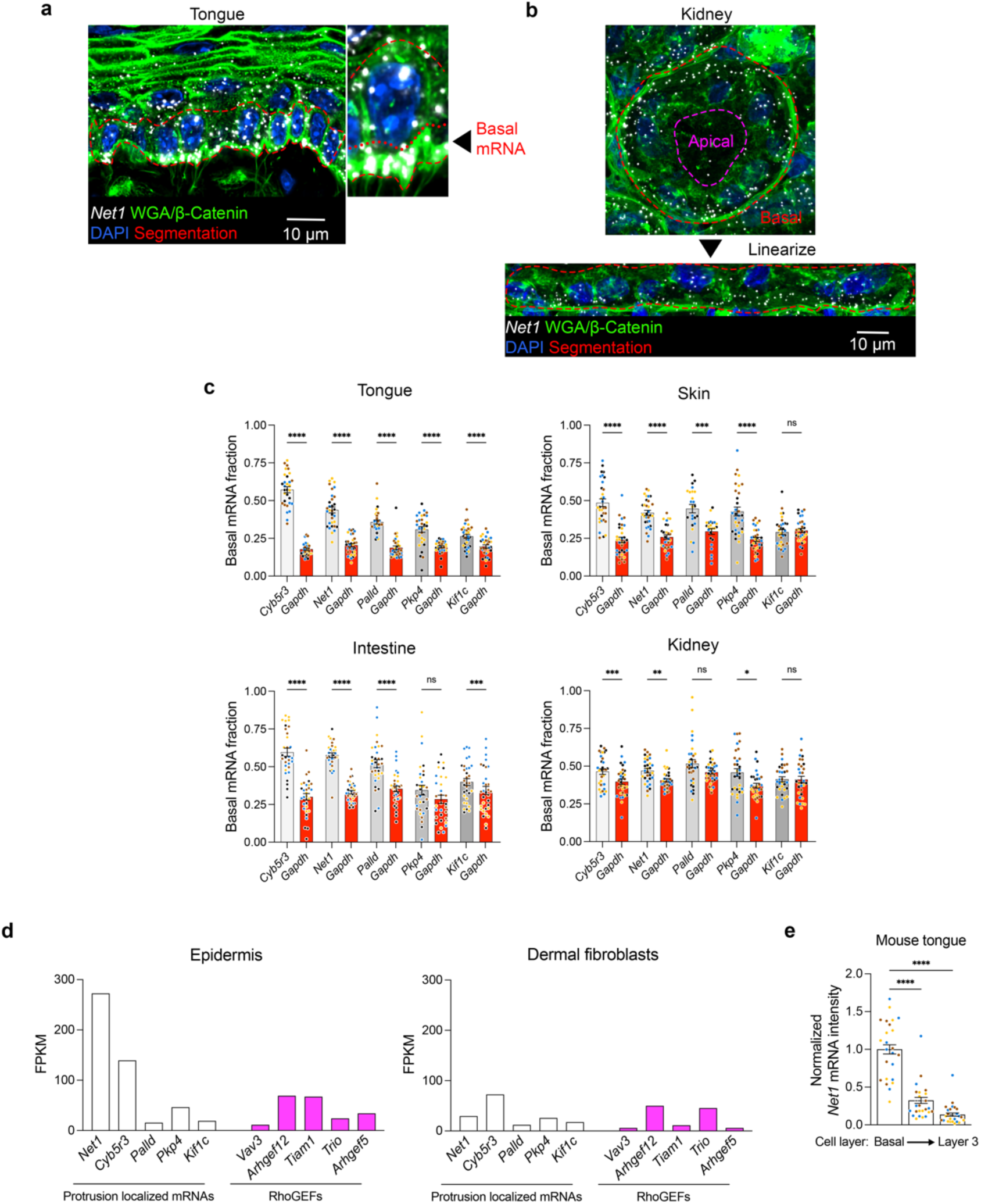
Protrusion localized mRNAs are basally localized across architecturally distinct tissues. **a**, Representative image of mouse tongue tissue stained for *Net1* mRNA and a combination of WGA and β-catenin. Manual segmentation of the basal cell layer is indicated by a red dashed line. The zoomed in inset highlights the position of the bottom 30% used to define the basal mRNA fraction. **b**, Representative tubular epithelial sheet from the kidney stained for *Net1* mRNA and a combination of WGA and β-catenin. For analysis, the image was first linearized and subsequently analyzed using the approach taken in stratified tissues. **c**, Basal mRNA fraction of several protrusion-localized mRNAs (*Cyb5r3*, *Net1*, *Palld*, *Pkp4*, and *Kif1c*; grey bars) and of the *Gapdh* control mRNA (red bars) in mouse tongue, skin, intestine, and kidney. **d**, Relative fluorescent intensity of *Net1* mRNA in tongue keratinocytes from the 1^st^ (basal), 2^nd^, and 3^rd^ epithelail layers. **e**, Protrusion-localized and RhoGEF mRNA expression in the skin epidermis vs. dermal fibroblasts derived from published datasets. *Net1* expression is notably higher in the epidermis to the point where it qualifies as a signature gene for the basal keratinocytes of the skin. Bar graph data are measurements from individual ROIs with mean ± SEM; n = 27-32 ROIs from N = 3-4 mice. Data from individual mice are represented with different colors. * P < 0.05, ** P < 0.01, *** P < 0.001, **** P < 0.0001 by repeated measures ANOVA followed by Sidak’s multiple comparisons test (c) or Kruskal-Wallis ANOVA followed by Dunn’s multiple comparisons test (d).

**Extended Data Fig. 2:**
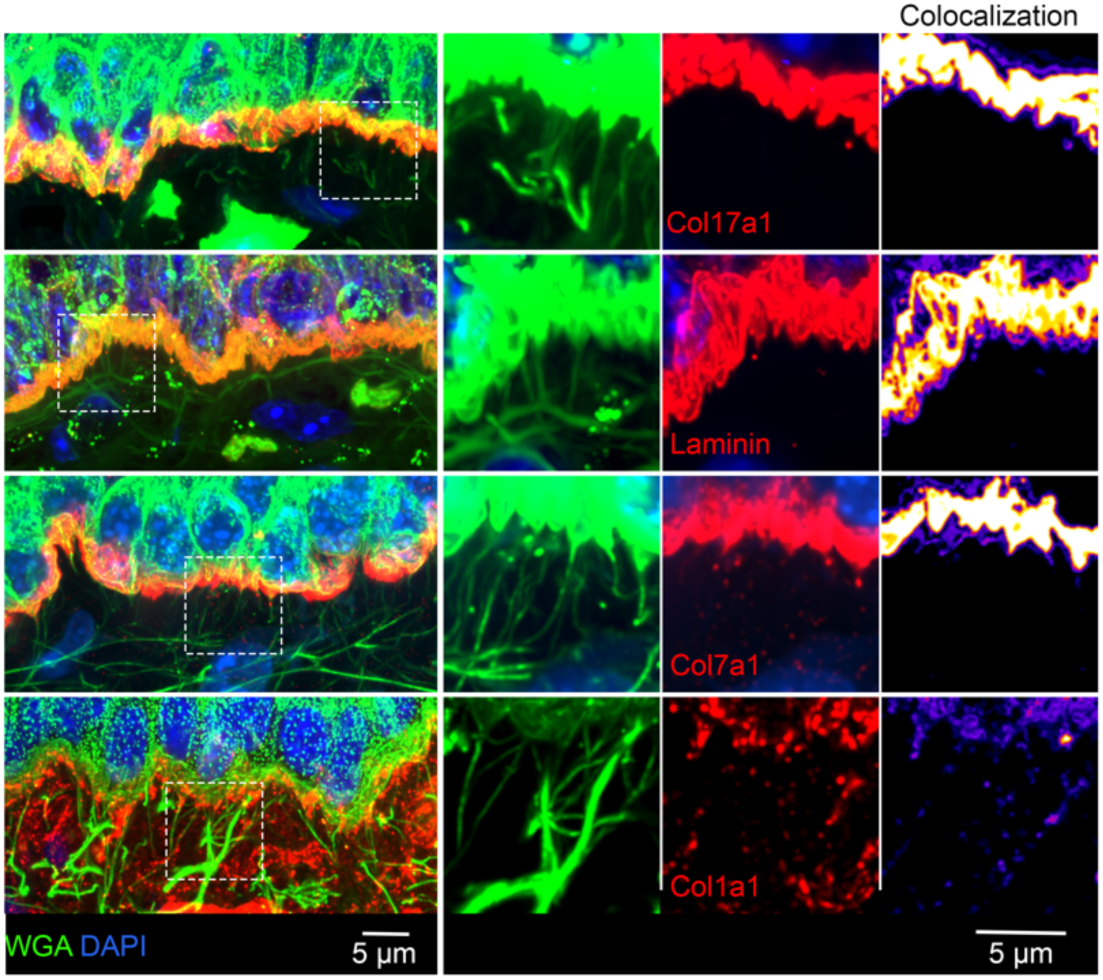
DEJ fibers are not comprised of hemidesmosome, basement membrane, or anchoring components. Representative images of mouse tongue tissue stained with WGA as well as with a panel of DEJ antibodies against Col17a1, pan-laminin, Col7a1, or Col1a1. Left panels: channel overlay images. Middle panels: individual channels (WGA:green; DEJ components: red) for the magnified boxed regions. Colocalization panels show pixel intensity overlap between WGA and DEJ channels. There was modest co-localization between DEJ fibers and Col1a1. Other DEJ components overlap with WGA at the region of the plasma membrane but not with WGA+ fibers below the DEJ.

**Extended Data Fig. 3:**
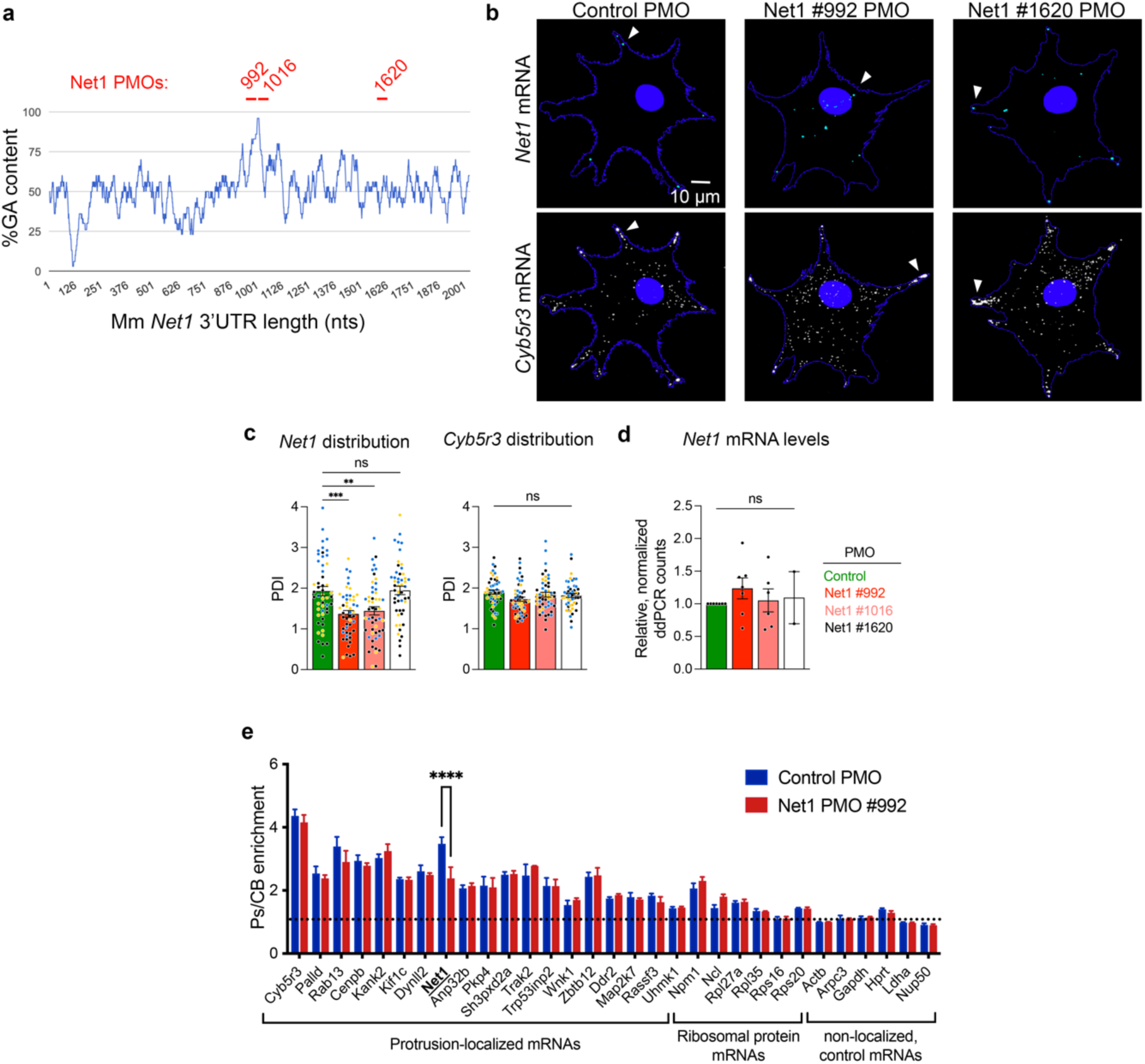
PMOs targeting the Mm *Net1* GA-rich region specifically alter *Net1* mRNA localization. **a**, GA content in 30nt windows along the *Net1* 3’UTR. Experimental Net1 PMOs were designed to target the *Net1* GA-rich region (#992 and #1016) or a different region (#1620) as a control (red lines indicate PMO locations). **b**, Representative images of *Net1* and *Cyb5r3* mRNA distribution in NIH/3T3 fibroblasts transfected with Control or Net1 targeting PMOs. Cell boundaries are indicated by a blue line. Arrowheads point to areas of RNA accumulation. Note that Net1 #992 PMO leads to perinuclear *Net1.* **c**, *Net1* and *Cyb5r3* mRNA distribution after PMO transfection as measured by a peripheral distribution index (PDI). PDI is an intensity weighted measure of the distribution of an RNA population relative to the center of the nucleus. A value above 1 is more peripheral, 1 is diffuse, and a value below 1 is a more perinuclear mRNA. n = 51 cells in 3 independent biological replicates. Data from independent experiments are shown in different colors. **d**, ddPCR count of *Net1* mRNA normalized to a housekeeping gene. n=2-7 biological and technical replicates **e**, Protrusion (Ps) and cell body (CB) fractions were isolated from 3T3 cells extending protrusions through transwell pores. The indicated mRNAs were detected by nanostring analysis to calculate a Ps/CB enrichment ratio. n=3. Bar graph data are individual measurements with mean ± SEM. ** P < 0.01, *** P < 0.001, **** P < 0.0001, ns: non-significant by ordinary one-way ANOVA followed by Dunnett’s multiple comparison test (c), mixed effects analysis with Dunnett’s multiple comparison test (d), or 2-way ANOVA followed by Sidak’s multiple comparison test (e).

**Extended Data Fig. 4:**
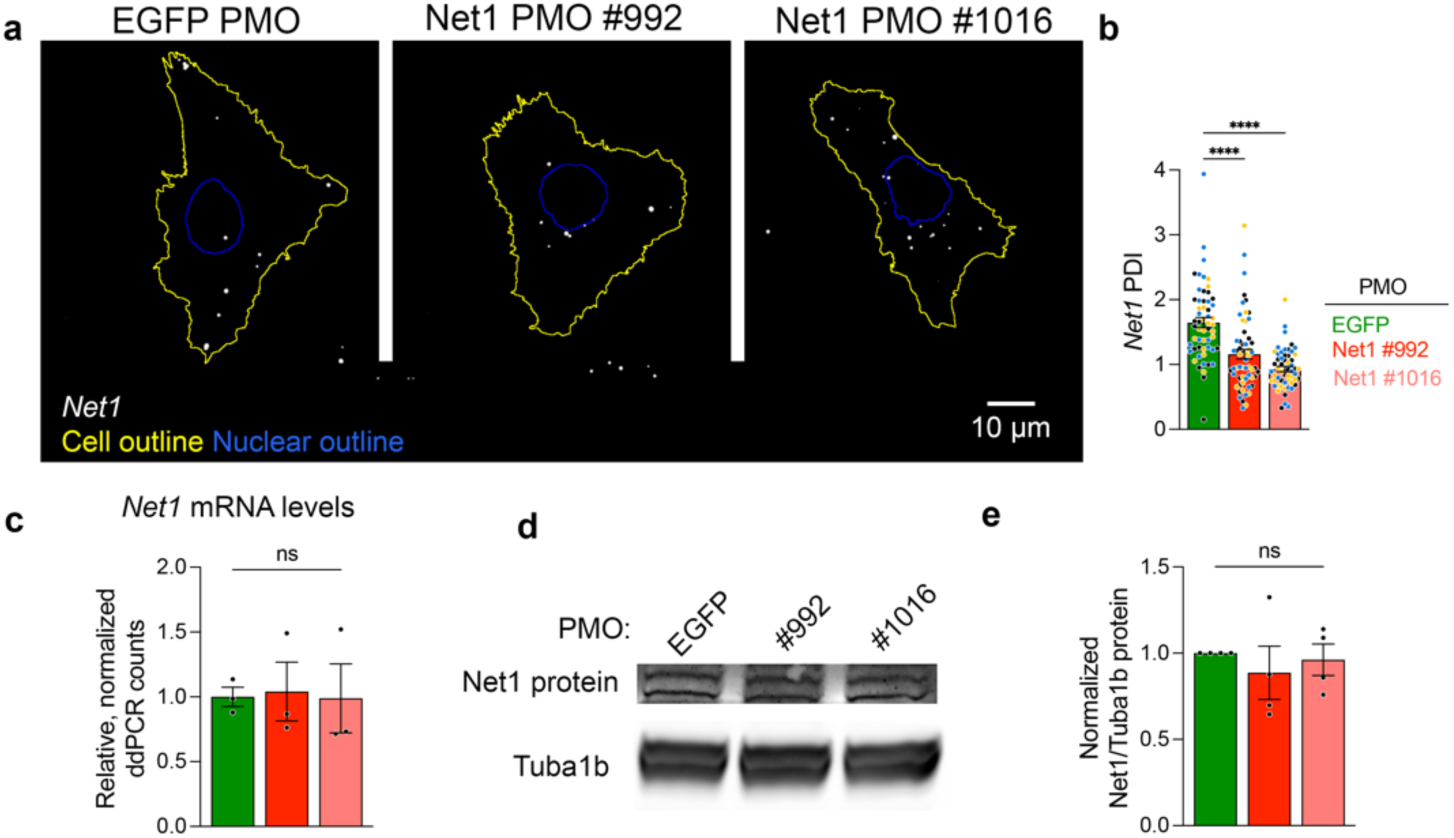
*Net1* PMOs alter *Net1* mRNA localization without affecting mRNA or protein abundance in mouse keratinocytes *in vitro*. **a**, Representative images of *Net1* mRNA localization in immortalized mouse keratinocytes treated with PMOs targeting *EGFP* or *Net1* mRNAs. **b**, Peripheral distribution of *Net1* measured by PDI in PMO treated cells. n = 56 cells in 3 independent biological replicates. Data from independent experiments are shown in different colors. **c**, ddPCR count of *Net1* mRNA normalized to a housekeeping gene *GusB*. n=3. **d**, Representative images of western blot detecting Net1 and Tuba1b protein from whole keratinocyte lysates. **e**, Quantification of Net1 protein normalized to Tuba1b. n=4. Bar graph data are individual measurements with mean ± SEM. **** P < 0.0001, ns: non-significant by Kruskal-Wallis ANOVA followed by Dunn’s multiple comparison test (b), or ordinary one-way ANOVA followed by Dunnett’s multiple comparison test (c,e).

**Extended Data Fig. 5:**
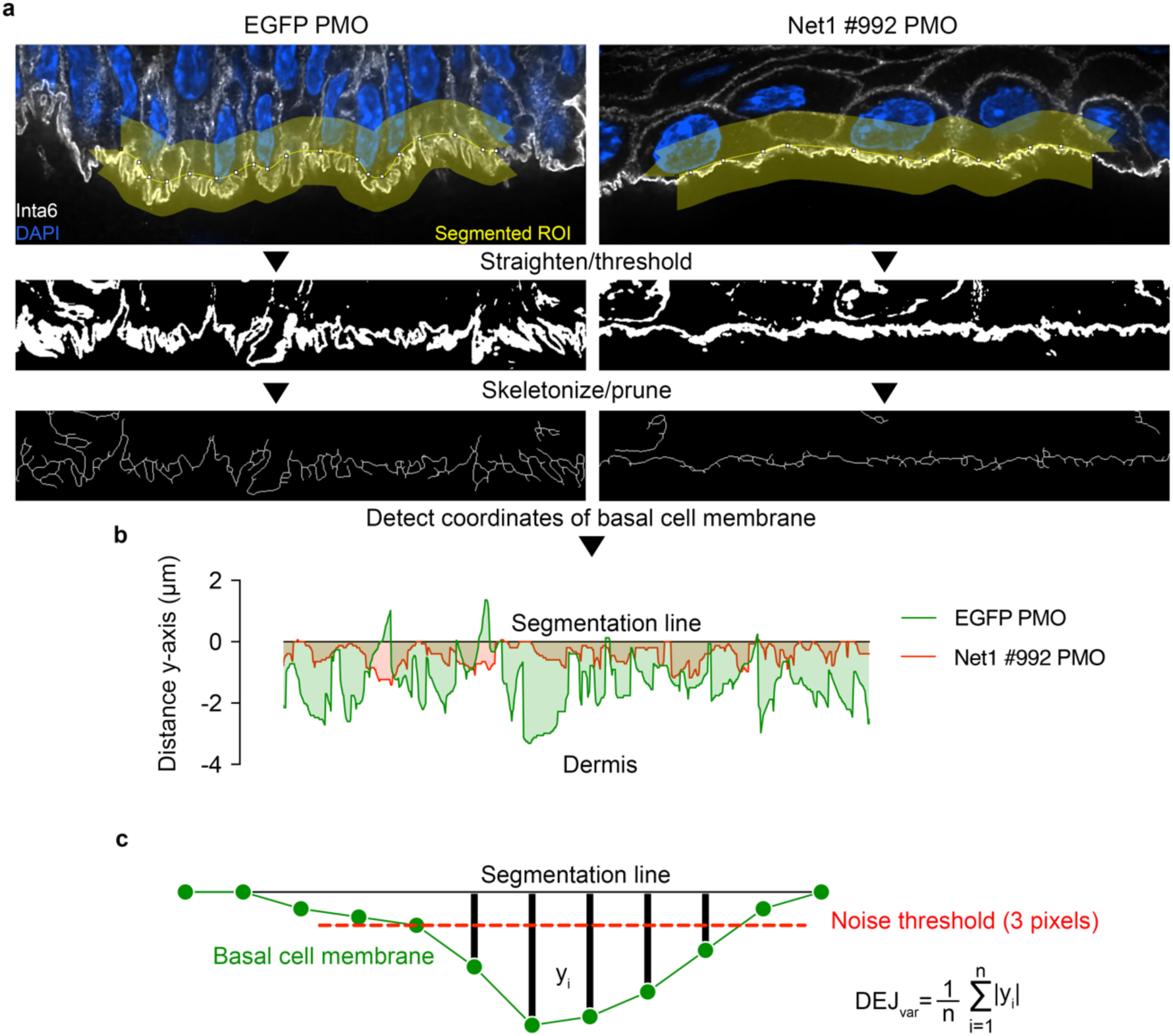
Analysis pipeline for measuring membrane topographical variation in the DEJ. **a**, For analysis, individual optical slices from tissue sections stained for Itga6 were used. Regions of the DEJ were isolated by drawing a segmented line ROI (120 pixel (7.8 μm) wide) centered on the basal cell membrane. The upper limits of protrusion-like structures were used as a reference. The ROI was digitally straightened and thresholded followed by skeletonization and pruning of incomplete membrane segments. **b**, From the mask in (**a**) the most basal coordinates in each x-position were derived. These basal coordinates were used to estimate y-axis variation relative to the original segmentation line. The graphical representation depicts x-position coordinates for the representative images from EGFP or Net1 PMO treated tissue. **c**, DEJ variation (DEJ_var_) was calculated as the average distance between the basal cell membrane coordinates (described in **b**) and the segmentation line. To limit the degree to which segmentation variability can influence the result a noise threshold of 3 pixels (∼.2 μm) was applied so that vertical displacement less than 3 pixels were set to 0. Multiple optical slices and segmentation lines were used to get a variability estimate across at least 100 μm of the DEJ for a single ROI, at least 5 ROIs were measured per animal.

**Extended Data Fig. 6:**
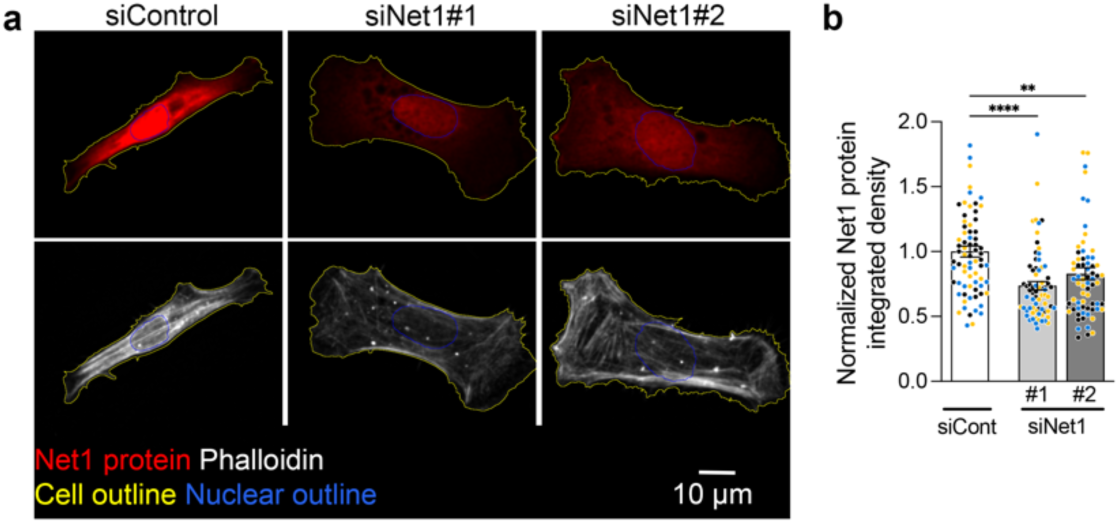
Net1 antibody validation for immunofluorescence in mouse keratinocytes. **a**, Representative Net1 protein immunofluorescence counterstained with phalloidin to visualize the actin cytoskeleton. Cells were treated with either a control or two different Net1 targeting siRNAs. Note: Net1 protein is predominantly nuclear in many *in vitro* cultured cells. **b**, Relative immunofluorescence intensity of Net1 protein. n = 22-28 cells per condition, N = 3 independent experiments. ** P < 0.01, **** P < 0.0001 by Kruskal-Wallis ANOVA followed by Dunn’s multiple comparison test.

**Extended Data Fig. 7:**
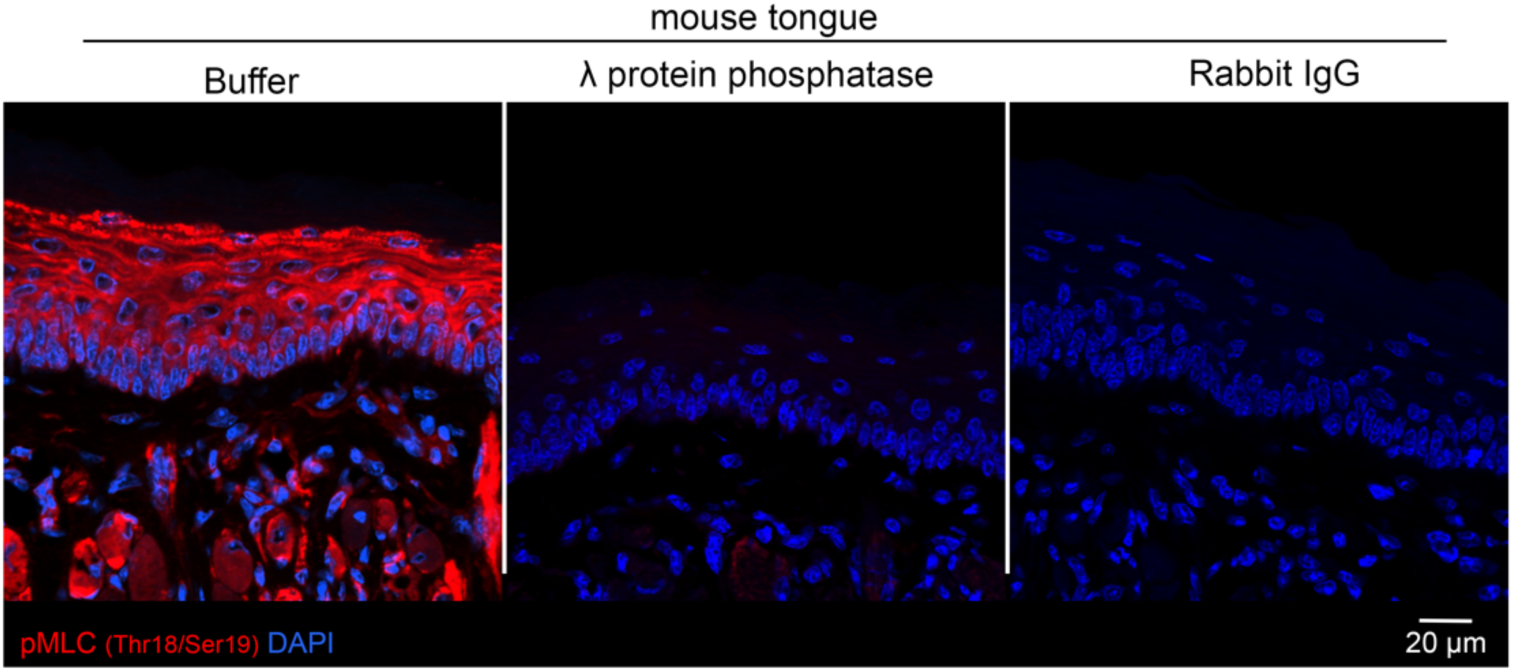
Myosin light chain (Thr18/Ser19) phosphorylation can be detected *in vivo* by immunofluorescence. Fixed mouse tongue sections were treated with λ protein phosphatase or buffer then stained with pMLC (Thr18/Ser19) primary antibody or Rabbit IgG. Immunofluorescent detection using an anti-rabbit secondary antibody indicated that pMLC primary antibody binding is diminished in tissues pre-treated with a serine, threonine, and tyrosine phosphatase.

**Supplementary Video 1: *Net1* mRNA and DEJ component staining in mouse tongue**. Representative serial optical sections of mouse tongue tissue. Upper panels: The basal cell membrane is visualized with the basal cell membrane marker Itga6, with or without overlay with *Net1* mRNA signal. Bottom panels: Visualization of the basal cell membrane and dermal fibers through WGA staining, with or without overlay with *Net1* mRNA signal. ∼3 μm thick serial optical sections were acquired with a ∼.065 μm step size.

**Supplementary Video 2: *Net1* mRNA and DEJ component staining in mouse tongue**. An additional representative serial optical sectioning of mouse tongue tissue. Upper panels: The basal cell membrane is visualized with the basal cell membrane marker Itga6, with or without overlay with *Net1* mRNA signal. Bottom panels: Visualization of the basal cell membrane and dermal fibers through WGA staining, with or without overlay with *Net1* mRNA signal. ∼3 μm thick serial optical sections were acquired with a ∼.065 μm step size.

**Extended Data Table 1: PMO and siRNA sequences used *in vitro* and *in vivo***

**Extended Data Table 2: Antibodies used for immunofluorescence**

**Extended Data Table 3: Probes for visualizing RNA**

## Notes

### Competing Interest Statement

The authors have declared no competing interest.

